# Indigenous probiotics *Lactobacillus reuteri* and *Enterococcus faecium* exhibit positive growth performance and disease prevention against extended-spectrum cephalosporin and fluoroquinolones resistant *Salmonella enterica* in broiler chicks

**DOI:** 10.1101/2023.07.10.548416

**Authors:** Abubakar Siddique, Roomana Ali, Amjad Ali, Saadia Andleeb, Nimatullah, Mudassar Mohiuddin, Samina Akbar, Muhammad Imran, Emily Van Syoc, Min Yue, Abdur Rahman, Erika Ganda

## Abstract

The rapid increase in antibiotic resistance poses a global threat to public health, necessitating the development of effective antimicrobial alternatives. This study compared an indigenous probiotic mix containing *Lactobacillus reuteri* and two strains of *Enterococcus faecium* to a commercial probiotic blend Protexin^R^ on the growth performance, mortality rate, histomorphology, serum immunoglobulins, and intestinal microflora of broiler chickens challenged with two multi drug resistant *Salmonella* serovars, Typhimurium and Enteritidis. Two hundred and forty day-old broiler chicks were randomly assigned to six treatment groups for 4 weeks: the treatment groups were; birds continuously supplemented with only indigenous probiotic strains (10^8^ CFU/mL) (IPRO-); birds challenged with *Salmonella* serovars 10^6^ (CFU/mL) (PC+); birds continuously supplemented with indigenous probiotic strains and challenged with *Salmonella* serovars (IPRO+); birds supplemented with Protexin^R^ and challenged with *Salmonella* serovars (CM+); birds supplemented with only Protexin^R^ (CM-); and birds with no *Salmonella* challenge or probiotics (negative control; PC-). The results revealed that IPRO- diets significantly improved feed conversion ratio (FCR) and increased body weight (BW) (*P* ≤ 0.05). No effect of probiotic treatments was observed on IPRO- and CM- on relative organ weights as compared to the negative control (PC-). The *Salmonella*- challenged group PC+ had the highest (20%) mortality rate and lowest BW. The IPRO- had significantly lower FCR (1.55) compared to PC- (1.86) and PC+ (1.95). The broilers in the IPRO- group showed significantly increased serum concentrations of IgA and IgG relative to both control groups (*P* ≤ 0.05). Morphological analysis of the ileum revealed significant increases (*P* ≤ 0.05) in the villus height and villus height/crypt depth in birds fed IPRO- compared with the PC+. Cecal *Lactobacillus* and *Enterococcus* counts were the highest (*P* ≤ 0.05) and *Salmonella* counts were the lowest (*P* ≤ 0.05) in the IPRO- group compared to the *Salmonella* infected group PC+. These results indicated that indigenous probiotic strains *Lactobacillus reuteri* and *Enterococcus faecium* can be an effective and low-cost alternative compared to commercial probiotics in the Pakistan poultry industry.

## INTRODUCTION

Salmonellosis is a foodborne infection caused by *Salmonella*, a gram-negative intestinal bacterium with significant public health significance in developing countries. (1). Consumption of contaminated meat or eggs from *Salmonella*-positive chickens is the primary source of human *Salmonella* infections (2). Up to 9% of fecal samples from poultry can be positive for *Salmonella* (3). *Salmonella enterica* serotypes Typhimurium, and Enteritidis are the most common *Salmonella* serotypes associated with human Salmonellosis (4). These serotypes are transferred to chickens vertically from parents or horizontally via environmental factors (5). *S*. Typhimurium and *S*. Enteritidis pose a threat to human health and can also be fatal to young chickens (6).

Antibiotics have long been employed in poultry production to treat and control infections, including Salmonellosis (7). However, the emergence of antimicrobial resistance (AMR) has become a global concern (8). Specifically, *Salmonella* infections have shown significant resistance to first-line antibiotics, including fluoroquinolones and cephalosporins, in clinical and veterinary practice (9). *Salmonella* serovars Typhimurium and Enteritidis have reportedly shown resistance to enrofloxacin, norfloxacin, ciprofloxacin, ceftriaxone, and cefepime (10, 11). Since antibiotics have largely been phased out of animal production, there is a critical need to identify safe and effective alternatives to control *Salmonella* infections.

Probiotics are live microorganisms that benefit the host animal (12). Probiotics have been widely tested as antibiotic alternatives with promising results in livestock as well as in humans. Indigenous probiotics are bacterial strains cultured from the species of interest and fed back as a probiotic, and these have been explored as a cost-effective alternative to commercially available probiotics. Indigenous probiotics have been tested to treat enteric infections, chronic inflammatory and allergic illnesses, immunomodulation and immunological stimulation, enhanced digestibility, and nutrient absorption in their host (13). The regular addition of indigenous probiotics in poultry diets may minimize the risk of infection with pathogens such as *Salmonella enterica*, *Campylobacter*, *E. coli*, and *Listeria monocytogenes*, and improve growth performance in birds (14, 15). Several reports have emerged on the successful use of indigenous or commercial probiotic strains to control or treat poultry infections including Salmonellosis, Colibacillosis, and necrotic enteritis (16–18). However, indigenous, and commercial probiotics have not been tested for the control of fluoroquinolones and extended spectrum cephalosporin resistant *Salmonella* serovars Typhimurium and Enteritidis so far. In the context of antibiotic resistance, it is critical to examine probiotic use to control multi-drug resistant (MDR) *Salmonella* strains in poultry.

In our previous study, we screened potential probiotic candidates evaluating their mucin adhesion properties and antagonistic activity against MDR *Salmonella enterica* strains (19). On the basis of these results, we identified three indigenous probiotic strains that we hypothesize could improve growth performance and immune function, modulate intestinal microflora, and control *Salmonella* serovars Typhimurium and Enteritidis infection in broiler chickens. Therefore, this study aimed to investigate the effect of an indigenous probiotic mixture (one *Lactobacillus reuteri* strain and two *Enterococcus faecium* strains) and a commercial probiotic mixture (Protexin^R^) on the growth performance, histomorphology, serum immunoglobulins and intestinal microflora of broiler chickens challenged with fluoroquinolones and extended-spectrum cephalosporin-resistant *Salmonella* serovars Typhimurium and Enteritidis.

## MATERIAL AND METHODS

### Ethical approval

The animal trial was approved by the Institutional Review Board of the Atta Ur Rahman School of Applied Biosciences, National University of Sciences and Technology, (Ref: No: IRB-132). The chickens were routinely monitored by a veterinarian.

### Animal husbandry

A total of 240 day-old Hubbard broiler chicks provided by a commercial hatchery (Sadiq Poultry Pvt Limited, Rawalpindi) with an average weight of 40g were randomized into six experimental treatments that lasted for four weeks (**Figure 1**). Each treatment consisted of four replicate pens containing ten birds in each pen. Each experimental group was housed in a floor pen with wood shavings as litter. The birds were fed an antibiotic-free basal feed comprised of corn, wheat, and soybean meal formulated according to NRC (1994) specifications. The starter diet was fed from 0-15 days; then, the grower-finisher diet was fed until the study ended (4 weeks). Feed analysis was carried out at Pakistan Council of Scientific and Industrial Research (PCSIR) laboratories in Lahore, Pakistan, and are shown in (**Table 1**). Lighting was provided for 18 hours/day throughout the experiment except from day 1 to day 7 when the lighting was provided for 23 hours. The room temperature was gradually reduced from 32°C during brooding to 26°C, and climate control was maintained. Feed and water were available *ad libitum* and bird performance was assessed weekly by recording the group weight and feed intake for each pen. The chickens were vaccinated against Newcastle diseases (ND.), avian pox, and Ghumbhuru on day 1 (20).

**Fig. 1:**
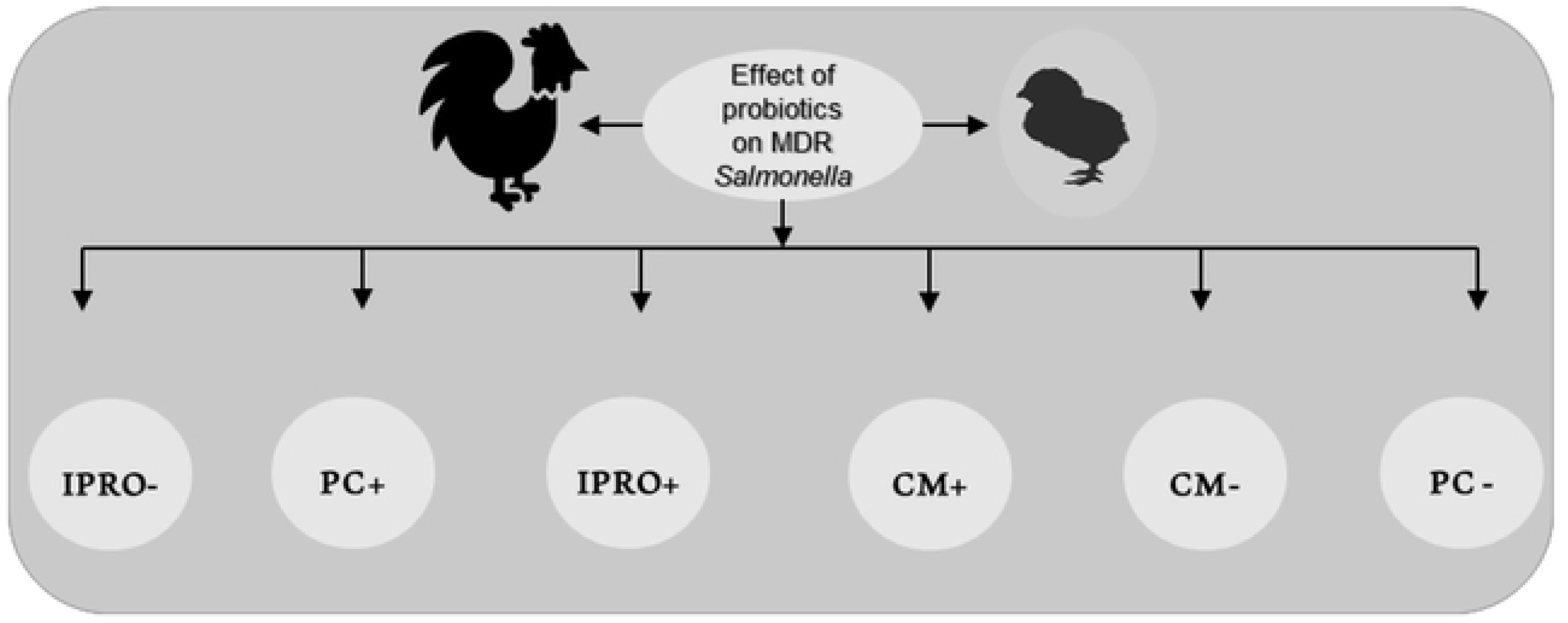
Six treatments in the study. IPRO-; Birds supplemented with only indigenous probiotic strains: PC+; Birds challenged with *Salmonella* serovars: IPRO+; Birds supplemented with indigenous probiotic strains and challenged with *Salmonella* serovars: CM+; Birds supplemented with Protexin^R^ and challenged with *Salmonella* serovars: CM-; Birds supplemented with Protexin^R^; PC-: Birds fed on normal diet with no probiotic supplementation or *Salmonella* serovar challenge.

**Table 1:**
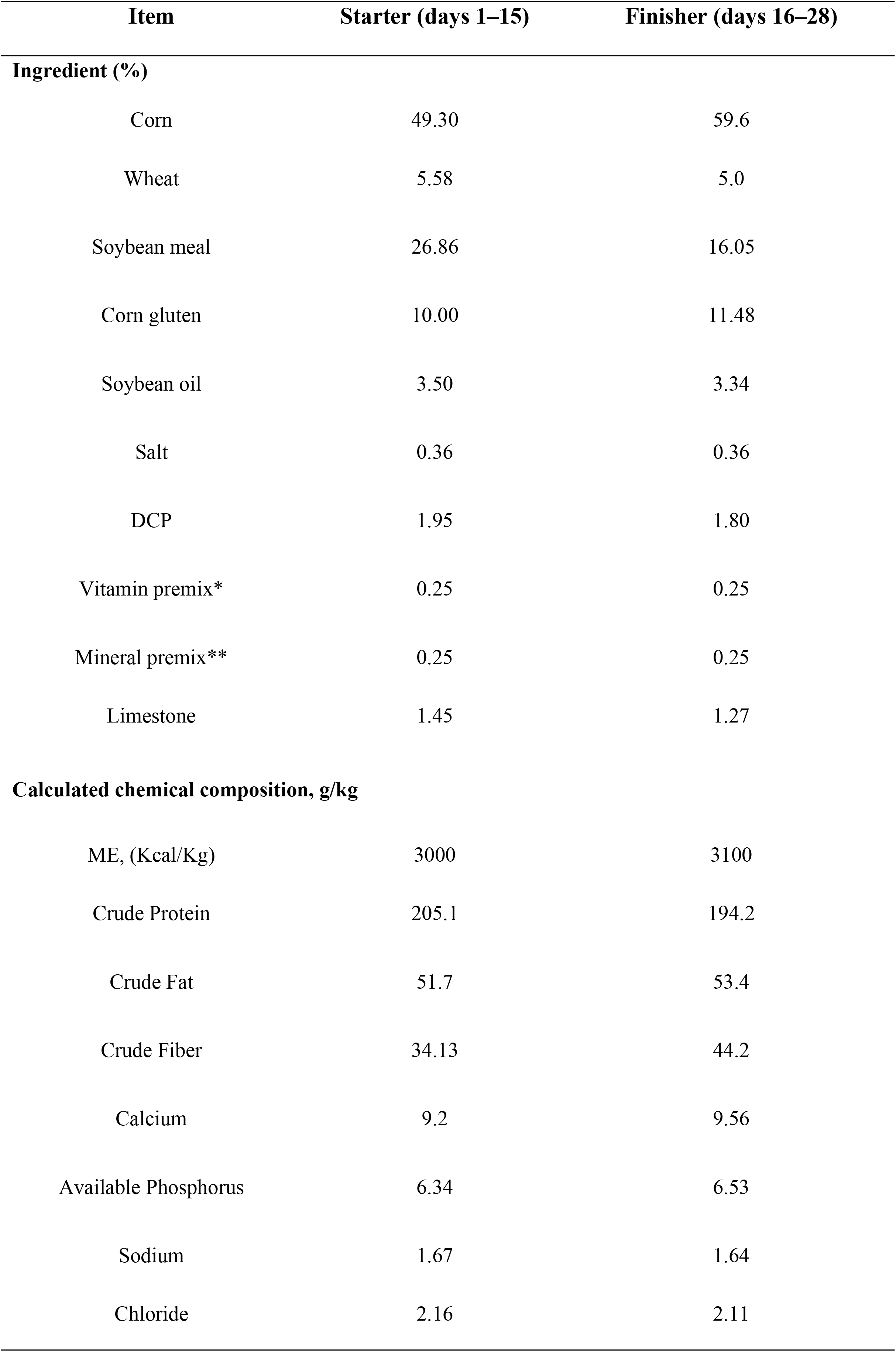
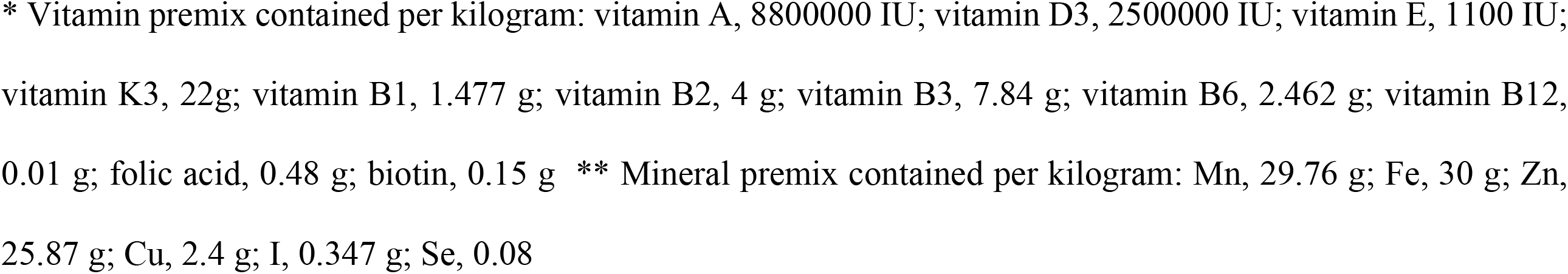
Composition of experimental diets used for broilers in the study.

| Item | Starter (days 1–15) | Finisher (days 16–28) |
| --- | --- | --- |
| **Ingredient (%)** |  |  |
| Corn | 49.30 | 59.6 |
| Wheat | 5.58 | 5.0 |
| Soybean meal | 26.86 | 16.05 |
| Corn gluten | 10.00 | 11.48 |
| Soybean oil | 3.50 | 3.34 |
| Salt | 0.36 | 0.36 |
| DCP | 1.95 | 1.80 |
| Vitamin premix* | 0.25 | 0.25 |
| Mineral premix** | 0.25 | 0.25 |
| Limestone | 1.45 | 1.27 |
| **Calculated chemical composition, g/kg** |  |  |
| ME, (Kcal/Kg) | 3000 | 3100 |
| Crude Protein | 205.1 | 194.2 |
| Crude Fat | 51.7 | 53.4 |
| Crude Fiber | 34.13 | 44.2 |
| Calcium | 9.2 | 9.56 |
| Available Phosphorus | 6.34 | 6.53 |
| Sodium | 1.67 | 1.64 |
| Chloride | 2.16 | 2.11 |

\* Vitamin premix contained per kilogram: vitamin A, 8800000 IU; vitamin D3, 2500000 IU; vitamin E, 1100 IU; vitamin K3, 22g; vitamin B1, 1.477 g; vitamin B2, 4 g; vitamin B3, 7.84 g; vitamin B6, 2.462 g; vitamin B12, 0.01 g; folic acid, 0.48 g; biotin, 0.15 g \*\* Mineral premix contained per kilogram: Mn, 29.76 g; Fe, 30 g; Zn, 25.87 g; Cu, 2.4 g; I, 0.347 g; Se, 0.08

### Probiotic strains

Three potential probiotic strains were previously isolated from chicken intestines and identified using 16S rRNA sequencing as *Lactobacillus reuteri* PFS4, *Enterococcus faecium* PFS13, and *Enterococcus faecium* PFS14. The indigenous probiotics were subcultured in Tryptic soy broth (TSB) and incubated at 37°C for 24 hours. The final concentration of each bacterial suspension of indigenous probiotics was adjusted to 0.5 McFarland standard (approx. 8 logs CFU/mL) concentrations by spectrophotometry (610 nm) (Biochrome Libra S22) followed by viable counting on tryptic soy agar (21).

The selected commercial probiotic mix was Protexin^R^ (provided by Hilton Pharma Pvt. Ltd., imported from the U.K.). Each dose (1mL) of this preparation contained 8 logs CFU/g of *Lactobacillus acidophilus, L. delbrueckii* subsp. bulgaricus, *L. plantarum, L. rhamnosus, Bifidobacterium bifidum, Enterococcus faecium, Streptococcus salivarius subsp. thermophiles, Aspergillus oryzae,* and *Candida pintolopesii*.

Both probiotic types were administered in drinking water. The commercial strain was administered as directed by the manufacturer. The indigenous probiotic mix was administered by adding at 8 log CFU/mL in drinking water. Both probiotics were administered in drinking water on the same day the chicks arrived.

### Pathogen challenge

*S.* Typhimurium FML15 and *S*. Enteritidis FML18, previously isolated from poultry feces, were selected as test pathogens. Both serovars were multi-drug-resistant with resistance to third-generation cephalosporin and quinolone (22). The serovars were subculture in TSB and incubated for 24 hours at 37°C. Pathogens were separated using centrifugation (three times at 5,000 RPM for 20 minutes) and washed in phosphate-buffered saline (PBS) (23). On days 5, 12, 19, and 26 of the trial, each bird in the experimental challenged groups was orally inoculated with 1 mL of 10^6^ CFU/mL of each serovar as previously described (24). Birds were examined at 12 h intervals for typical clinical manifestations of Salmonellosis until the end of the experiment. Clinical signs and mortality were recorded accordingly (2).

### Growth performance and carcass traits

Growth performance parameters, including body weight gain (BWG), feed intake (FI), and feed conversion ratio (FCR; feed intake/weight gain), were calculated at the end of every week and over the experimental period (28 days). The chicks were observed for clinical signs, morbidity, and mortality (mortality rate (%) = the number of dead chicks per group/number of chicks per group × 100%). Mortality was recorded daily from each pen as it occurred, and feed was adjusted according to mortality (25).

Broilers were euthanized with a 0.5mL/kg intraperitoneal injection of 6.3% sodium pentobarbital (Sigma Aldrich, USA). On day 7 and day 28 three chickens from each replicate pen in control and experimental groups were randomly selected, weighed, and euthanized to determine relative organ weight. Before euthanasia, the birds fasted for 12 h. The weights of the gizzard, ileum, cecum, liver, bursa, spleen, and kidney were calculated as a relative percentage of live body weight. Small intestine pH was quantified by measuring diluted intestinal digesta in phosphate buffer saline with a pH meter (26).

### Microbiological studies

On days 7, 14, 21, and 28, three birds from each replicate were euthanized, and 1-2 grams of fresh caecum digesta and liver contents were immediately collected within one hour of dissection for microbial enumeration. 1 g of each sample was serially diluted from 10^−1^ to 10^−10^ in PBS for the enumeration of *Salmonella enterica* serovars, *Lactobacillus* spp., and *Enterococcus* spp. by conventional microbiological techniques using selective media including *Salmonella Shigella* agar with rifampicin 20µg/mL, MRS agar, and M17 agar with rifampicin 60µg/mL, respectively (27). Bacterial enumerations were conducted in duplicate, and the mean was expressed as log_10_ CFU/g (base-10 logarithm colony-forming units per gram) of cecal digesta and liver tissues.

### Serum IgA and IgG content

On day 7 and 28, three chickens from each replicate were randomly selected and blood samples were collected into 5-mL vacuum tubes, for the analysis of serum IgA and IgG level using an ELISA kit (Thermo Fisher Scientific, USA) according to (28). 96-well microtitration plates (Wuxi NEST Biotechnology Co., Ltd, USA) were coated with 100 μL of antigen diluted in 0.1 M carbonate buffer and incubated overnight at 4°C. The plates were washed three times with PBS-Tween 20. To prevent nonspecific binding, wells were blocked with 100 μL of 8% nonfat dry milk PBS-Tween 20 and incubated for 1.5 h at 37°C. For IgA analysis, 100 μL of 1:200 dilution of the serum in 8% nonfat dry milk-PBS-Tween 20 were added to the plates in duplicates and incubated for 1.5 h at room temperature. For IgG analysis, 100 μL of 1:200 dilution of the serum in 8% nonfat dry milk-PBS-Tween 20 were added to the plates in duplicates and incubated for 1.5 h at room temperature. The optical density was read at 450 nm using a microplate ELISA reader. IgA and IgG values are reported as the mean optical density.

### Ileal morphometric and histological analysis

On day 7 and 28, one bird close to the average BW from each pen was euthanized, and a 2 cm section of ileum was collected and stained with hematoxylin and eosin (H&E) (29). The sections were examined at 100x magnification and photographed using a microscope coupled with a camera (CX21; Olympus Optical Co., Ltd, Tokyo, Japan). Using ImageJ software, three villi per cross-section were measured to determine the villus height, crypt depth, and the ratio of villus height to crypt depth (30).

### Statistical analysis

The overall differences between different treatments were analyzed by one-way ANOVA using the SPSS Statistics 22.0 (SPSS Inc., Chicago, IL) statistical software. Differences between the treatment groups were considered statistically different at *P* ≤ 0.05. When the interaction effects were significant (*P* ≤ 0.05), differences between means of different treatment were analyzed by Tukey’s least-square test.

## RESULTS

### Clinical symptoms, mortality, and gross lesions

Clinical symptoms were observed in all groups challenged with *S*. Enteritidis and *S.* Typhimurium (PC+, IPRO+, and CM+) (**Fig. 1A- 1E Supplementary figure**). Clinical symptoms were observed in the PC+ treatment within a few hours of the inoculation; chickens huddled in the corners of the partition showed drowsiness, loss of appetite, and inhibition in drinking. They were generally depressed and hesitant to move, and some of the chicks developed thin, yellowish diarrhea. The clinical symptoms seen in the birds were transient and gradually dissipated within 16-24 hours, and complete recovery in the infected birds occurred within 48 hours.

Mortality differed by probiotic treatment in the infected groups (**Table 2**). From day 1 to 5, each group showed normal performance, but from day 6 to 28, the mortality rate in the PC+ group was significantly higher (20%) (*P* ≤ 0.05) than in the other groups. During the trial, no mortality was observed in the IPRO- group.

**Table 2.** Overall broiler body weight gain (BWG), feed intake (FI), feed conversion ratio (FCR) and mortality % for 28 days.

| Items | Treatments<sup>1</sup> |  |  |  |  |  | Statistics |  |
| --- | --- | --- | --- | --- | --- | --- | --- | --- |
|  | I<sup>1</sup>PRO- | PC+ | I<sup>1</sup>PRO+ | CM+ | CM- | PC- | SEM* | P-Value |
| ABW (g) | 995<sup>a</sup> | 770<sup>c</sup> | 900<sup>b</sup> | 890<sup>b</sup> | 900<sup>b</sup> | 885<sup>b</sup> | 29.62 | 0.01 |
| AFI (g) | 1,550<sup>b</sup> | 1,500<sup>c</sup> | 1,530<sup>ab</sup> | 1,570<sup>c</sup> | 1,550<sup>b</sup> | 1,600<sup>a</sup> | 18.75 | 0.002 |
| FCR | 1.55<sup>d</sup> | 1.95<sup>a</sup> | 1.68<sup>b</sup> | 1.76<sup>b</sup> | 1.67<sup>b</sup> | 1.86<sup>ab</sup> | 0.058 | 0.01 |
| Mortality (%) | 0.0 | 20.0 | 2.5 | 7.5 | 2.5 | 0.0 |  |  |

\*Standard error of mean (SEM), <sup>a-c</sup>Means with different superscripts within the same row differ significantly ( $P \leq 0.05$ ).
<sup>1</sup>I<sup>1</sup>PRO-; Birds supplemented only indigenous probiotic strains; CM+; Birds challenged with *Salmonella* serovars: I<sup>1</sup>PRO+; Birds supplemented with indigenous probiotic strains and challenged with *Salmonella* serovars; CM+; Birds supplemented with Protexin<sup>R</sup> and challenged with *Salmonella* serovars; CM-; Birds supplemented with only Protexin<sup>R</sup> ; PC+: Birds fed on normal diet

### Growth performance

The weekly progress of broiler growth performance during the whole experiment is shown in (**Figures 2A and 2B**). Average body weight (BW) of day-old chicks was 40.3 ± 0.87 g and did not differ among treatments. The mean body weight during the experiment was significantly higher (*P* ≤ 0.05) for IPRO- (995g) compared with CM- (900g), PC+ (770g) and PC- (885g) in (Figure 2A). Feed intake during 1^st^ and 2^nd^ week was not significantly differed among different treatments. It was significantly differed among treatments during week 3 and 4 (Figure 2B). In week 3 and week 4, IPRO- (833g and 1550g, respectively) had significantly higher FI than PC+ (805g and 1500g). The feed conversion ratio (FCR) was significantly lower in IPRO- (1.55) than CM- (1.67), PC- (1.86), and PC+ (1.95). The improved FCR (*P* ≤ 0.05) in IPRO+ (1.68) and CM+ (1.76) compared to PC+ (1.95) may be attributed to the improvement in clinical symptoms after probiotics administration.

**Figure 2.**
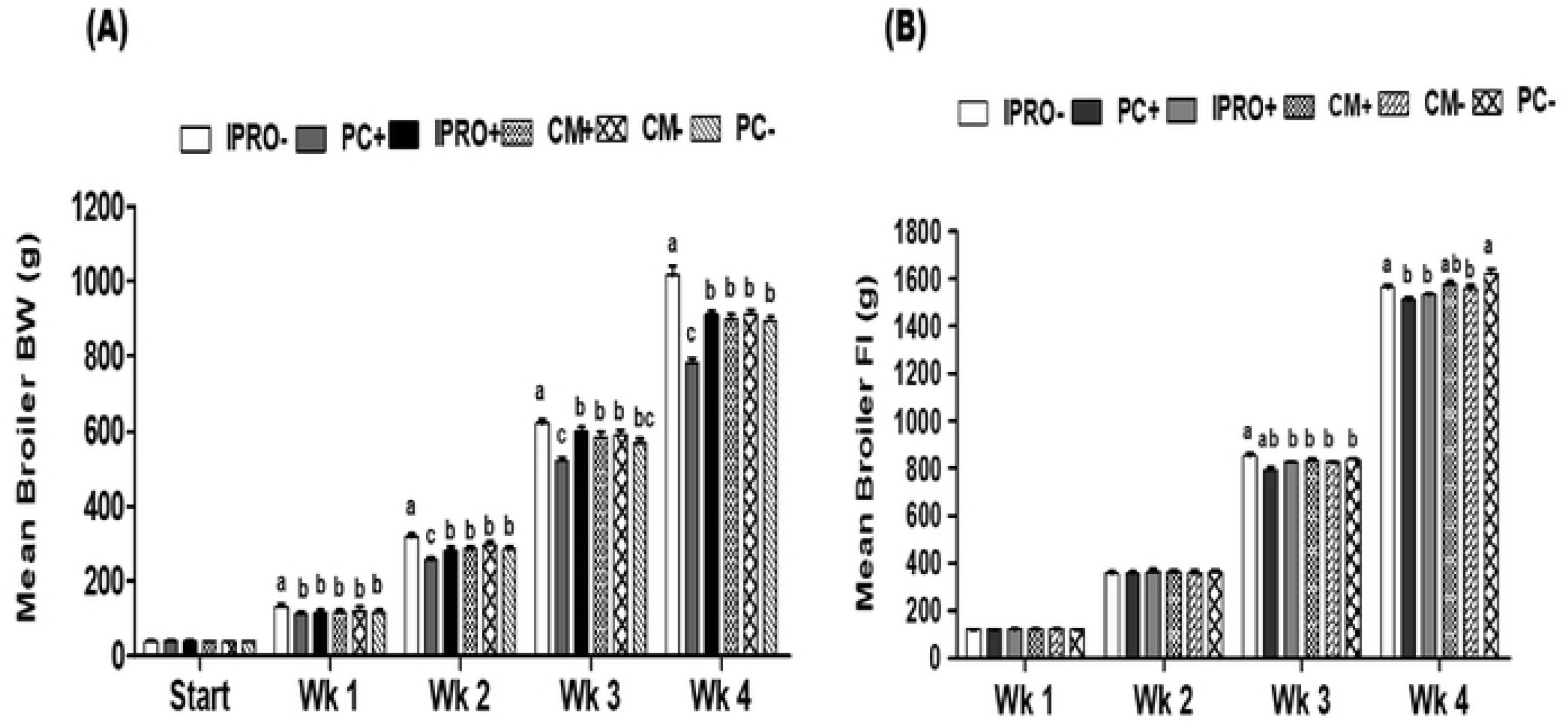
(A) Broiler weekly body weight (BW) and (B) Broiler weekly feed intake (FI) in IPRO-; Birds supplemented with only indigenous probiotic strains: CM+; Birds challenged with *Salmonella* serovars: IPRO+; Birds fed on indigenous probiotic strains and challenged with *Salmonella* serovars: CM+; Birds supplemented with Protexin^R^ and challenged with *Salmonella* serovars: CM-; Birds supplemented with only Protexin^R^ ; PC+: Birds fed on normal diet. Bars represent means for the 3 replicates per treatment ± SEM. Within the same week, bars with different letters (a,b and c) differ significantly (P ≤ 0.05).

### Organ weights and intestinal pH

The means of organ weights relative to the BW and intestinal pH are shown in (**Table 3**). On day 7, the relative weights of the liver, spleen, caecum, and small intestine were higher (*P* ≤ 0.05) in the S*almonella*-infected (PC+) than in the other treatment groups. No significant changes (*P* ≤ 0.05) in the weight of any other organ were detected in birds after the *Salmonella* infection on day 7. Except for the liver and small intestine, there was no significant difference (*P* ≤ 0.05) in the relative weight of organs between the challenged and non-challenged groups after day 28. In addition, the pH of the intestine remained unaffected by dietary supplementations.

**Table 3:** Effects of probiotic supplementation and *Salmonella* challenge on relative organ weights and intestinal pH (shown as % body weight) of broilers at days 7 and 28.

| Item | Treatment |  |  |  |  |  | SEM | *P-value* |
| --- | --- | --- | --- | --- | --- | --- | --- | --- |
|  | IPro- | PC+ | IPro+ | CM+ | CM- | PC- |  |  |
| Day 7 |  |  |  |  |  |  |  |  |
| Liver | 4.0<sup>b</sup> | 4.15<sup>a</sup> | 4.03<sup>b</sup> | 4.04<sup>b</sup> | 4.01<sup>b</sup> | 4.02<sup>b</sup> | 0.022 | 0.004 |
| Spleen | 0.17<sup>b</sup> | 0.22<sup>a</sup> | 0.17<sup>b</sup> | 0.18<sup>b</sup> | 0.16<sup>b</sup> | 0.17<sup>b</sup> | 0.008 | 0.001 |
| Pancreas | 0.79 | 0.81 | 0.76 | 0.77 | 0.76 | 0.73 | 0.018 | .41 |
| Gizzard | 4.91 | 5.0 | 5.01 | 5.07 | 4.9 | 4.89 | 0.173 | 0.12 |
| Small intestine | 8.84<sup>b</sup> | 9.52<sup>a</sup> | 8.8<sup>b</sup> | 8.85<sup>b</sup> | 8.76<sup>b</sup> | 8.81<sup>b</sup> | 0.189 | <0.0001 |
| Intestinal pH | 6.62 | 6.74 | 6.69 | 6.71 | 6.67 | 6.68 | 0.19 | 0.23 |
| Colon | 0.61 | 0.64 | 0.63 | 0.62 | 0.61 | 0.61 | 0.012 | 0.36 |
| Caecum | 2.08<sup>b</sup> | 2.15<sup>a</sup> | 2.09<sup>b</sup> | 2.04<sup>b</sup> | 2.07<sup>b</sup> | 2.08<sup>b</sup> | 0.014 | 0.0005 |
| Bursa | 0.43 | 0.45 | 0.44 | 0.46 | 0.43 | 0.45 | 0.004 | 0.22 |
| Day 28 |  |  |  |  |  |  |  |  |
| Liver | 2.97<sup>b</sup> | 3.3<sup>a</sup> | 3.1<sup>b</sup> | 3.1<sup>b</sup> | 3.1<sup>b</sup> | 2.96<sup>b</sup> | 0.060 | 0.01 |
| Spleen | 0.13 | 0.15 | 0.14 | 0.14 | 0.12 | 0.13 | 0.004 | 0.81 |
| Pancreas | 0.35 | 0.37 | 0.35 | 0.36 | 0.34 | 0.37 | 0.007 | 0.39 |
| Gizzard | 3.15 | 3.18 | 3.16 | 3.17 | 3.14 | 3.18 | 0.008 | 0.69 |
| Small intestine | 5.68<sup>b</sup> | 5.97<sup>a</sup> | 5.77<sup>b</sup> | 5.74<sup>b</sup> | 5.71<sup>b</sup> | 5.76<sup>b</sup> | 0.035 | 0.001 |
| IntestinalpH | 6.54 | 6.59 | 6.62 | 6.51 | 6.63 | 6.61 | 0.25 | 0.13 |
| Colon | 0.28 | 0.29 | 0.27 | 0.28 | 0.28 | 0.28 | 0.003 | 0.17 |
| Caecum | 0.62 | 0.64 | 0.64 | 0.63 | 0.63 | 0.62 | 0.003 | 0.34 |
| Bursa | 0.25 | 0.27 | 0.26 | 0.24 | 0.26 | 0.26 | 0.004 | 0.43 |

<sup>a-b</sup>Means within a row lacking a common superscript are significantly different ( $P \leq 0.05$ ).

### Histomorphology of the intestine

The results of morphological analysis of ileum and jejunum on days 7 and 28 are shown in (**Table 4**). Morphological study of the ileum revealed significant increases in villus height (*P* ≤ 0.05) and villus height/crypt depth ratio (*P* ≤ 0.05) as well as decreases in crypt depth (*P* ≤ 0.05) in IPRO- and CM-treatments as compared to the control group. However, no significant difference in jejunal epithelium height or tunica muscularis thickness was observed in all treatments.

**Table 4:** Ileal morphometric analysis of Broiler at day 7 and day 28

| Item | Dietary Treatment |  |  |  |  |  | SEM | *P-value* |
| --- | --- | --- | --- | --- | --- | --- | --- | --- |
|  | IPRO- | PC+ | IPRO+ | CM+ | CM- | PC- |  |  |
| Day 7 |  |  |  |  |  |  |  |  |
| Villus height, μm | 510<sup>a</sup> | 447<sup>c</sup> | 477<sup>b</sup> | 478<sup>b</sup> | 498<sup>a</sup> | 486<sup>b</sup> | 8.72 | 0.003 |
| Crypt depth, μm | 90<sup>b</sup> | 110<sup>a</sup> | 94<sup>b</sup> | 95<sup>b</sup> | 92<sup>b</sup> | 90<sup>b</sup> | 3.08 | 0.08 |
| Villi: Crypt | 5.7<sup>a</sup> | 4.06<sup>d</sup> | 5.07<sup>c</sup> | 5.04<sup>c</sup> | 5.41<sup>b</sup> | 5.4<sup>b</sup> | 0.23 | <0.001 |
| Epithelium height, μm | 74 | 76 | 71 | 72 | 69 | 79 | 2.08 | 0.19 |
| Thickness of tunica muscularis, μm | 159 | 164 | 163 | 163 | 162 | 161 | 0.73 | 0.16 |
| Day 28 |  |  |  |  |  |  |  |  |
| Villus height, μm | 795<sup>a</sup> | 695<sup>c</sup> | 745<sup>b</sup> | 734<sup>b</sup> | 785<sup>a</sup> | 787<sup>a</sup> | 19.08 | <0.0001 |
| Crypt depth, μm | 121<sup>c</sup> | 146<sup>a</sup> | 139<sup>a</sup> | 141<sup>a</sup> | 128<sup>b</sup> | 132<sup>b</sup> | 3.70 | 0.003 |
| Villi: Crypt | 6.57<sup>a</sup> | 4.77<sup>b</sup> | 5.28<sup>c</sup> | 5.2<sup>c</sup> | 5.98<sup>d</sup> | 5.89<sup>d</sup> | 0.29 | <0.0001 |
| Epithelium height, μm | 91 | 99 | 95 | 96 | 94 | 92 | 3.29 | 0.18 |
| Thickness of tunica muscularis, μm | 257 | 252 | 254 | 249 | 254 | 258 | 4.18 | 0.15 |

<sup>a-c</sup> means without a common superscript differ significantly within a row ( $P \leq 0.05$ )

### Serum IgA and IgG content analysis

The results of serum immunoglobulin IgA and IgG levels at day 7 and day 28 were shown in (**Figure 3a-3d**). The results showed that the level of IgA and IgG significantly increased from day 7 to day 28 in all treatments. When compared to the negative control (PC-) group, all treatments had significantly higher levels (*P* ≤ 0.05) of IgA. On day 7 and day 28, the level of IgA was highest in the indigenous probiotic group IPRO- (118.5 and 293.4 g/mL), but it was not significantly different from the commercially available probiotic (CM-) (111.8 and 282.3 g/mL). The level of IgG was higher (*P* ≤ 0.05) in IPRO- as compared to positive control PC+ and negative control PC-, whereas the other groups exhibited no differences on day 7 (**Fig 3c**). Moreover, at day 28, the level of IgG was significantly higher in IPRO- compared to other treatments.

**Figure 3:**
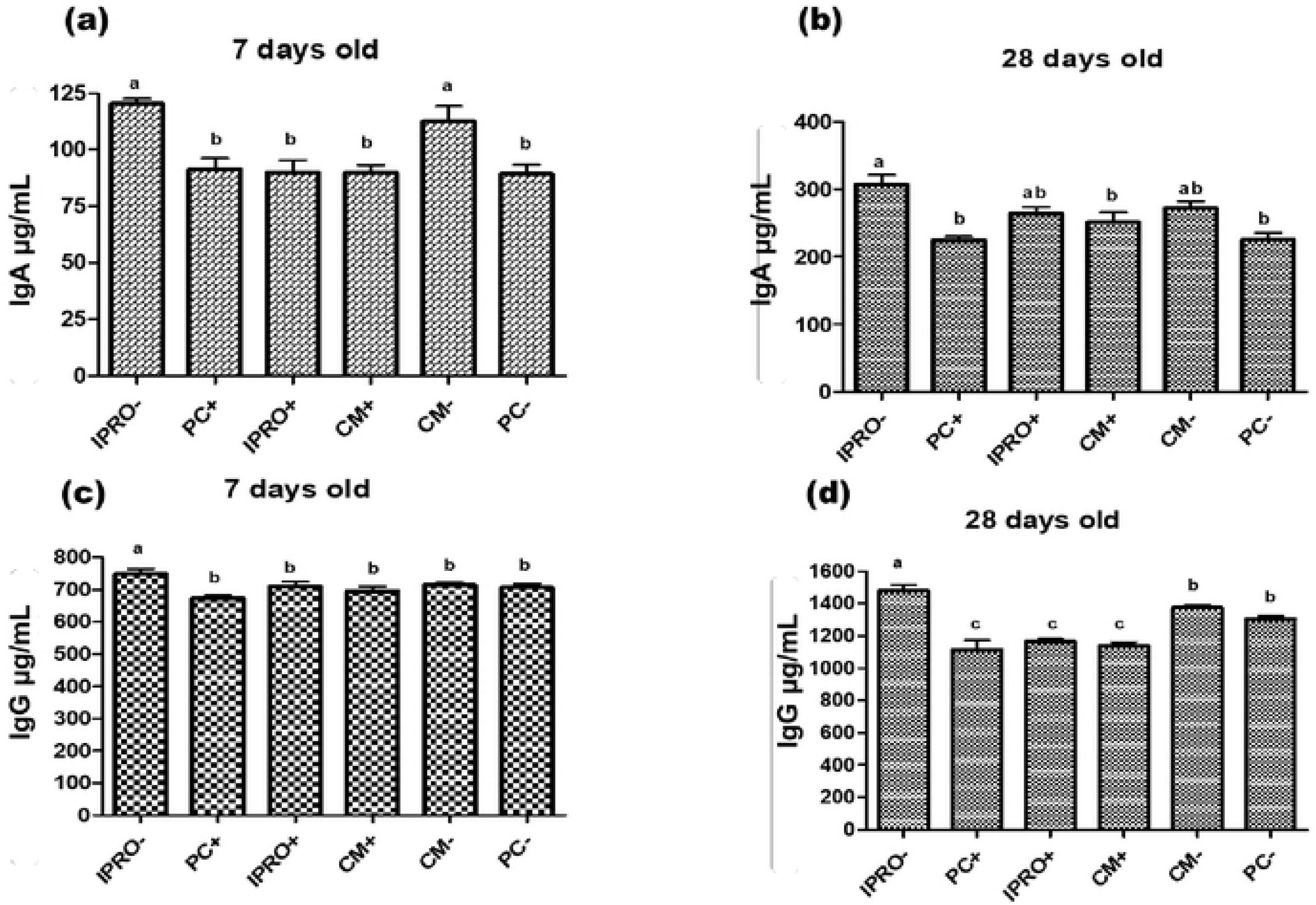
Impact of treatments on serum concentrations of IgA, and IgG in broilers. (a) The level of IgA in 7 days old. (b) The level of IgA in 28 days old. (c) The level of IgG in 7 days old. (d) The level of IgG in 28 days old. Bars represent means for the 3 replicates per treatment ± SEM. Within the same week, bars with different letters (a, b and c) differ significantly (P ≤ 0.05).

### Bacteriological analysis

The effect of indigenous and commercial probiotics on the *Salmonella* loads in caecum and livers of all the challenged groups on days 7, 14, 21, and 28 are shown in (**Figure 4A-4B**). All cecal and liver samples tested negative for *Salmonella* on day 0 of the study. The cecal and liver *Salmonella* load was significantly lower (*P* ≤ 0.05) in IPRO+ and CM+ groups compared to the positively challenged group PC+ from day 7 to day 28. The effect of indigenous and commercial probiotics on the total *Lactobacillus* and *Enterococcus* count on days 7, 14, 21, and 28 are shown in (**Figure 5A-5B**). The *Salmonella* challenged group (PC+) had a significantly (*P* ≤ 0.05) lower total *Enterococcus* and *Lactobacillus* count than other treatments. The *Lactobacillus* and *Enterococcus* counts did not differ significantly between the indigenous and commercial probiotic groups.

**Figure 4:**
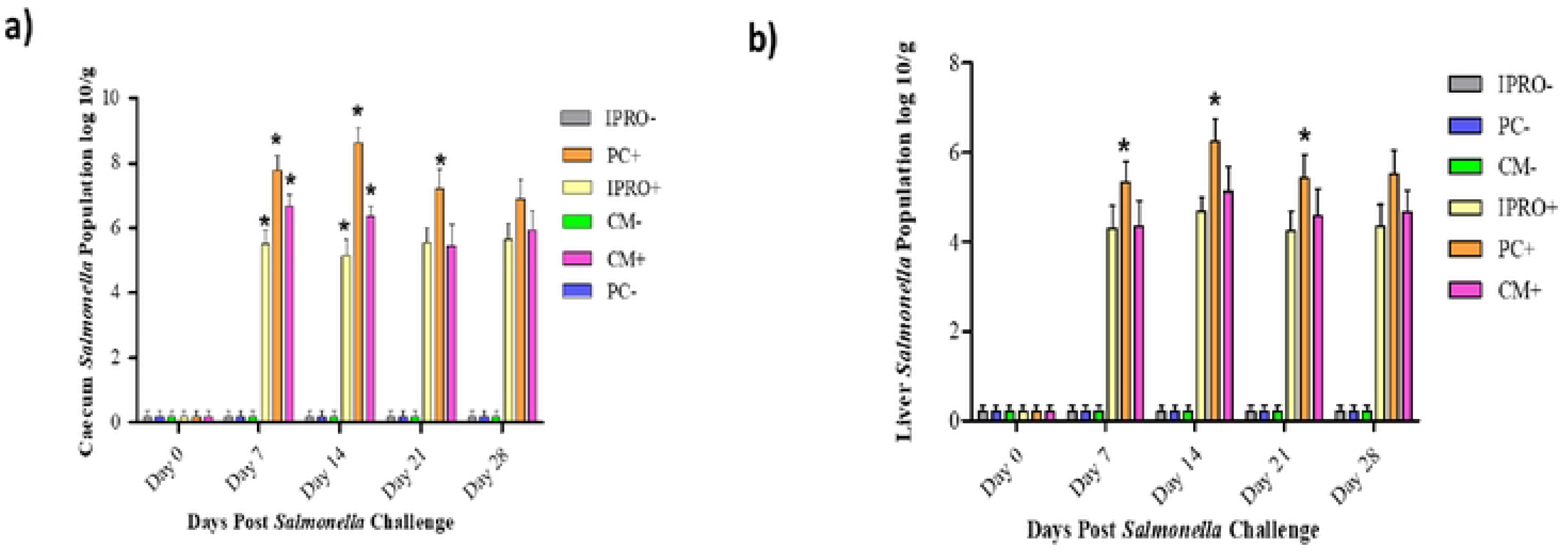
Salmonella sp. quantification. **A);** Effect of indigenous and commercial probiotic supplementation on cecal Salmonella cntel’ica scrovars cecal digesta load after Salmonella infection in broile,· birds. **B);** Effect of indigenous and commercial probiotic supplementation on hepatic Salmonella enterica serovars liver load after Salmonella infcc1ion in broiler birds. P,·obiotics were supplemented **in** water from the day of hatch until 28 d of age. At 0, 7, **14,** 21 and 28 d after infection, cecal contents and liver were analyzed for S. cntcrica scrovars by plate count method, collected and expressed as IOlog values. Bars (±SEM) with • superscript differ significantly’ (P ≤ 0.05).

**Figure 5:**
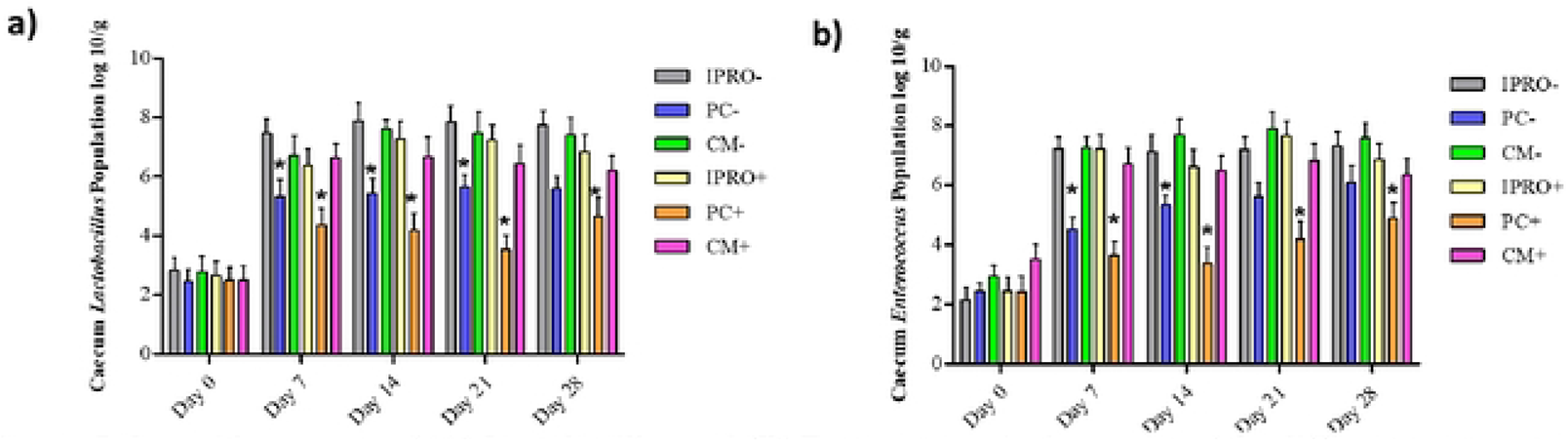
Logarithm coun1s of **(A)** Lactobacillus and (R) En1crococcus in the cecum of 1hc diffcrcn1 trcaimcnt groups of chicks. Probiolics were supplemented in waler from lhe day of halch uniii 28 d of age. At 0, 7, 14, 21and 28 d cecal con1en1s were analyzed for L.ac1obacillusand En1erococcus by plale counl method. Values with* differed significantly within a bird trial (P ≤ 0.05). All data are in log IO CFU/g contents and expressed as (±SEM).

### Histological Examinations

The histopathological changes in the small intestine from various groups at day 28 are presented in (**Figure 6A-6F**). The intestinal villi in the positive challenged group (PC+) were the shortest and arranged irregularly. The *Salmonella* challenge in the treatment group with commercial probiotic mix (CM+) caused partial enteritis in the intestinal villi, characterized by loss of tissue architecture and accumulation of tissue debris, mucus, and desquamated intestinal sheets in the lumens (Figure 6d). The height and width of the intestinal villi were more regular in the indigenous probiotic group (IPRO+) than in the positive control group (PC+). The villi height and width appeared normal in the indigenous probiotic group (IPRO+), however, there was partial coagulated necrosis in the lumen and separation of some villi tips observed in (IPRO+).

**Figure 6:**
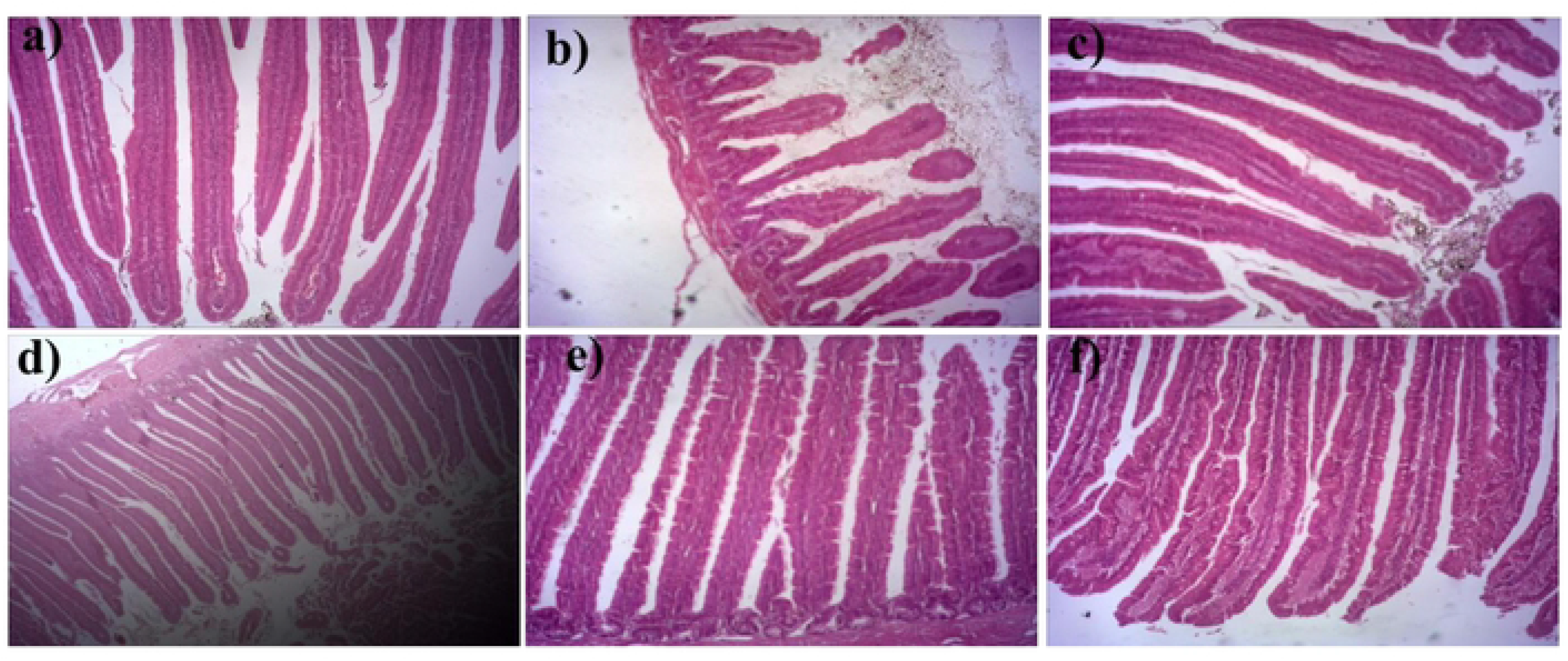
Histopathologic changes of the ileum in different treatment groups of chicks**. (a)** = IPRO-; **(b)** = PC+; (**c)** = IPRO+; (**d)** = CM+ (**e)** = CM-; (**f)** = PC-. Magnification is 40X.

## DISCUSSION

The intestinal microbiota plays an important role in broiler growth and provide resistance against pathogen invasion. Indigenous probiotics that are isolated from the normal flora of the species, are of particular interest as a cost-effective alternative to commercial probiotic mixtures. Some indigenous potential probiotics have been associated with improved health and performance, product quality, and pathogen resistance (31). Indigenous and commercial probiotic strains are being tested to prevent *Salmonella enterica* infections in poultry production as an alternative to antimicrobials (15, 32). In this study, all chicks were healthy before infection with *Salmonella* serovars and the highest mortality rate (20.0%) after *Salmonella* infection was observed in *Salmonella* positive challenged group with no probiotic supplementation (PC+). The clinical symptoms and death rate after infection were consistent with previous findings (33). This indicates that the model of Salmonellosis in chicks was successfully established.

Broiler performance parameters BWG and FCR were significantly improved in indigenous probiotic treatment (IPRO-) compared with control groups (PC+) and (PC-) and commercial probiotic treatment group (CM-), demonstrating a beneficial effect of indigenous probiotics compared to commercial strains. The exact mechanism by which indigenous strains showed better growth results than a commercial probiotic is unknown, but the probiotic source may be one reason for this variation (34). The commercial probiotics used in our study were derived from an unknown host, whereas the indigenous bacteria were recovered from the intestines of poultry birds. The advantages of indigenous *L. reuteri* and *E. faecium* on broiler growth performance are consistent with other studies in broiler chickens using different indigenous probiotic strains such as *Lactobacillus*, *Bacillus*, and *Enterococcus* (15, 35, 36). This may result from beneficial effects of indigenous lactic acid bacterial (LAB) strains on intestinal microflora and morphology, improving nutrient digestibility and absorption (37). Birds infected with *Salmonella* had reduced BWG and feed conversion ratio (FCR). However, the addition of probiotics to the diet helps in the improvement of growth performance. Reduced growth performance in the *Salmonella* infected birds may be due to decreased feed intake. These results were consistent with previous findings (38, 39).

In our study, a significant increase was observed in the relative weight of the liver, spleen, caecum, and small intestine in the *Salmonella*-challenged group (PC+) on days 7 and 28 compared to all other treatment groups. These results were consistent with the previous findings of (40), who reported similar changes in the relative weight of organs in chickens infected with *Salmonella* strains. The addition of probiotics (indigenous or commercially prepared) had no significant effect on the relative weight of organs (gizzard, liver, pancreas, spleen, and bursa), which is also consistent with previous findings (41). The relative weights of the critical organs can often be used as an indicator of changes in the morphology of the gut (30).

Healthy intestinal morphology is essential to growth performance. Higher villus height and lower crypt depth improve nutrient absorption in the intestines by providing more surface area and a faster rate of nutrient absorption (42). This study showed that indigenous (IPRO-) as well as commercial probiotic (CM-) treatment groups significantly increased the villus height and crypt depth and increased the ratio of villus height to crypt depth as compared to control groups PC+ and PC-. The increase in the villi height and villus height to crypt depth ratio leads to the increase in the absorption of nutrients in the small intestine, which affects the production of birds (43). The ratio of villus height to crypt depth was significantly lower in *Salmonella*- infected birds (PC+) on days 7 and 28, demonstrating the negative impact of *S. enterica* infection on intestinal morphology. According to a previous study, probiotics have been reported to increase villus height in broiler chickens. (44).

Serum immunoglobulins are biomarkers of immune status in animals (45). Probiotics can enhance the humoral immune response by increasing immunoglobulins levels (26, 46). In our study, serum IgA and IgG levels of birds in IPRO- group were the highest compared to other treatments on days 7 and 28. According to a previous study, *Lactobacillus* mixed probiotics increased IgA levels and reduced mortality in *Salmonella*-infected birds (47). According to earlier findings, birds fed orally with *Lactobacillus acidophilus*, *Lactobacillus reuteri*, and *Lactobacillus salivarius* have a modified systemic humoral immune response (48, 49). Similar findings were observed in a previous study in which serum IgG and IgA levels in probiotic-supplemented groups were significantly higher than in controls (50).

Chickens infected with *S*. Enteritidis or *S*. Typhimurium may have an altered balance of intestinal flora (39, 51). *Lactobacillus* is well known for reducing pathogenic *Salmonella* colonization and regulating the equilibrium of the intestinal micro-ecosystem (52). *L. reuteri* and *E. faecium* strains colonize the digestive tracts of chicks and can improve their intestinal health (52, 53). Our findings indicated that supplementation with indigenous probiotics and a commercial probiotic combination in drinking water was associated with lowered *Salmonella* counts and increased *Lactobacillus* and *Enterococcus* counts in the caecum and liver. Because the intestinal flora of freshly hatched chicks is incomplete, early *Lactobacillus* consumption can aid in colonizing a healthy intestinal flora and increase the chick’s resistance to *Salmonella* infection that is not specific to the host (48). Previous findings have found that the intestinal flora balance is perturbed in chicks infected with *Salmonella* serovars (39). A previous study reported that the *E. faecium* EF55 reduced the cecal *Salmonella* in the broilers (54). Similarly, other studies have shown that probiotics, *L. reuteri,* and *E. faecium*, reduced the concentration of *Salmonella* strains in chicks (55, 56). The probiotic supplemented diet also increased the amount of total cecal *Lactobacillus* and total *Enterococcus*, according to our findings. Similarly, (57) demonstrated that a high dose of *E. faecium* stimulated other lactic acid bacteria in the small intestine. A previous study showed that a diet with the probiotic *L. reuteri* increased the lactic acid bacteria colonization in the ileum of chicks (58). The mucosa’s health and the intestinal tract’s microscopic structure can be good indicators of the intestinal tract’s response to enteric infections and the efficacy of bioactive feed supplements. According to our findings, birds treated with *Salmonella* (PC+) displayed symptoms of intestinal tissue injury, including degeneration and necrosis of intestinal villi, which is consistent with previous findings (59, 60). This variation in ileum histology could be related to *Salmonella’s* preference for the cecum as a target tissue.

## CONCLUSION

In conclusion, broilers fed with the indigenous probiotic strains (*L. reuteri* PFS4, *E. faecium* PFS13, and *E. faecium* PFS14) had increased weight gain and improved feed conversion ratio compared to a commercial probiotic mixture (Protexin) in *Salmonella* challenged and non-challenged groups. The indigenous probiotic strains effectively reduced *S.* Typhimurium and *S*. Enteritidis load in the caecum and liver, and increased *Lactobacillus* and *Enterococcus* abundance in caecum. They also improved serum immunoglobulins IgA and IgG, and villi height. The indigenous probiotic mixture can be an effective alternative to antibiotics by improving growth performance and protecting against multi-drug resistant *Salmonella* serovars.

## ACKNOWLEDGMENTS

The authors thank all Lab animal house staff of NUST for providing facilities and a feed preparation machine during the experiment.

## Author Contributions

All authors listed have made a substantial, direct, and intellectual contribution to the work and approved it for publication.

## Patent

A patent from this data has been submitted provisionally under patent application no.534/2022.

## REFERENCES

1. Pui C, Wong W, Chai L, Tunung R, Jeyaletchumi P, Hidayah N, et al. Salmonella: A foodborne pathogen. International Food Research Journal. 2011;18(2).

2. Pulido-Landínez M. Food safety-Salmonella update in broilers. Animal Feed Science Technology. 2019;250:53–8.

3. Fearnley E, Raupach J, Lagala F, Cameron S. Salmonella in chicken meat, eggs and humans; Adelaide, South Australia, 2008. International journal of food microbiology. 2011;146(3):219–27.

4. Zou M, Keelara S, Thakur SJ. Molecular characterization of Salmonella enterica serotype Enteritidis isolates from humans by antimicrobial resistance, virulence genes, and pulsed-field gel electrophoresis. Foodborne Pathogens Diseases. 2012;9(3):232–8.

5. Liljebjelke KA, Hofacre CL, Liu T, White DG, Ayers S, Young S, et al. Vertical and horizontal transmission of Salmonella within integrated broiler production system. Foodbourne Pathogens Diseases 2005;2(1):90–102.

6. Tang B, Siddique A, Jia C, Ed-Dra A, Wu J, Lin H, et al. Genome-based risk assessment for foodborne Salmonella enterica from food animals in China: A One Health perspective. International Journal of Food Microbiology. 2023;390:110120.

7. Vieco-Saiz N, Belguesmia Y, Raspoet R, Auclair E, Gancel F, Kempf I, et al. Benefits and inputs from lactic acid bacteria and their bacteriocins as alternatives to antibiotic growth promoters during food-animal production. Frontiers in microbiology. 2019;10:57.

8. Sartelli M, C Hardcastle T, Catena F, Chichom-Mefire A, Coccolini F, Dhingra S, et al. Antibiotic Use in Low and Middle-Income Countries and the Challenges of Antimicrobial Resistance in Surgery. Antibiotics. 2020;9(8):497.

9. Jajere SM. A review of Salmonella enterica with particular focus on the pathogenicity and virulence factors, host specificity and antimicrobial resistance including multidrug resistance. Veterinary world. 2019;12(4):504.

10. Ballesté Delpierre CC. Quinolone resistance acquisition and impact on virulence in Salmonella enterica: a cost-benefit matter: Universitat de Barcelona; 2015.

11. Hossain MS. Extended-spectrum β-lactamase producing and Quinolone resistance Salmonella spp. in Dhaka city retail meat: an emerging public health concern of Bangladesh: Brac University; 2018.

12. Sánchez B, Delgado S, Blanco-Míguez A, Lourenço A, Gueimonde M, Margolles A. Probiotics, gut microbiota, and their influence on host health and disease. Molecular nutrition food Research International. 2017;61(1):1600240.

13. Saad N, Delattre C, Urdaci M, Schmitter J-M, Bressollier P. An overview of the last advances in probiotic and prebiotic field. LWT-Food Science Technology. 2013;50(1):1–16.

14. Markowiak P, Śliżewska K. The role of probiotics, prebiotics and synbiotics in animal nutrition. Gut pathogens. 2018;10(1):1–20.

15. Salehizadeh M, Modarressi MH, Mousavi SN, Ebrahimi MT. Effects of probiotic lactic acid bacteria on growth performance, carcass characteristics, hematological indices, humoral immunity, and IGF-I gene expression in broiler chicken. Tropical animal health production. 2019;51(8):2279–86.

16. Botlhoko TD. Performance of Clostridium perfringens-challenged broilers inoculated with Effective Microorganisms: University of Pretoria; 2010.

17. Chambers JR, Gong J. The intestinal microbiota and its modulation for Salmonella control in chickens. Food Research International. 2011;44(10):3149–59.

18. Fancher CA, Zhang L, Kiess AS, Adhikari PA, Dinh TT, Sukumaran AT. Avian pathogenic Escherichia coli and Clostridium perfringens: Challenges in no antibiotics ever broiler production and potential solutions. Microorganisms. 2020;8(10):1533.

19. Siddique A, Azim S, Ali A, Adnan F, Arif M, Imran M, et al. Lactobacillus reuteri and Enterococcus faecium from Poultry Gut Reduce Mucin Adhesion and Biofilm Formation of Cephalosporin and Fluoroquinolone-Resistant Salmonella enterica. Animals. 2021;11(12):3435.

20. Olnood CG, Beski SS, Choct M, Iji PA. Novel probiotics: Their effects on growth performance, gut development, microbial community and activity of broiler chickens. Animal Nutrition. 2015;1(3):184–91.

21. Higgins S, Higgins J, Wolfenden A, Henderson S, Torres-Rodriguez A, Tellez G, et al. Evaluation of a Lactobacillus-based probiotic culture for the reduction of Salmonella enteritidis in neonatal broiler chicks. Poultry Science. 2008;87(1):27–31.

22. Siddique A, Azim S, Ali A, Andleeb S, Ahsan A, Imran M, et al. Antimicrobial Resistance Profiling of Biofilm Forming Non Typhoidal Salmonella enterica Isolates from Poultry and Its Associated Food Products from Pakistan. Antibiotics. 2021;10(7):785.

23. Silva I, Vellano I, Moraes A, Lee I, Alvarenga B, Milbradt E, et al. Evaluation of a Probiotic and a Competitive Exclusion Product Inoculated In Ovo on Broiler Chickens Challenged with Salmonella Heidelberg. Brazilian Journal of Poultry Science. 2017;19(1):19–26.

24. Olnood CG, Beski SS, Iji PA, Choct M. Delivery routes for probiotics: Effects on broiler performance, intestinal morphology and gut microflora. Animal Nutrition. 2015;1(3):192–202.

25. Gao J, Zhang H, Yu S, Wu S, Yoon I, Quigley J, et al. Effects of yeast culture in broiler diets on performance and immunomodulatory functions. Poultry Science. 2008;87(7):1377–84.

26. Zhang Z, Cho J, Kim I. Effects of Bacillus subtilis UBT-MO2 on growth performance, relative immune organ weight, gas concentration in excreta, and intestinal microbial shedding in broiler chickens. Livestock Science. 2013;155(2-3):343–7.

27. Engberg R, Hedemann M, Steenfeldt S, Jensen B. Influence of whole wheat and xylanase on broiler performance and microbial composition and activity in the digestive tract. Poultry science. 2004;83(6):925–38.

28. Sadeghi AA, Shawrang P, Shakorzadeh SJ. Immune response of Salmonella challenged broiler chickens fed diets containing Gallipro®, a Bacillus subtilis probiotic. Probiotics antimicrobial proteins. 2015;7(1):24–30.

29. Velleman S, Coy C, Anderson J, Patterson R, Nestor K. Effect of selection for growth rate and inheritance on posthatch muscle development in turkeys. Poultry science. 2003;82(9):1365–72.

30. Oxford JH, Selvaraj RK. Effects of glutamine supplementation on broiler performance and intestinal immune parameters during an experimental coccidiosis infection. Journal of Applied Poultry Research. 2019;28(4):1279–87.

31. Noohi N, Ebrahimipour G, Rohani M, Talebi M, Pourshafie M. Evaluation of potential probiotic characteristics and antibacterial effects of strains of Pediococcus species isolated from broiler chickens. British poultry science. 2016;57(3):317–23.

32. Wibisono FM, Wibisono FJ, Effendi MH, Plumeriastuti H, Hidayatullah AR, Hartadi EB, et al. A review of salmonellosis on poultry farms: public health importance. Systematic Reviews in Pharmacy. 2020;11(9):481–6.

33. Gast RK, Porter Jr RE. Salmonella infections. Diseases of poultry. 2020:717–53.

34. Abdollahi-Arpanahi D, Soltani E, Jafaryan H, Soltani M, Naderi-Samani M, Campa-Córdova AI. Efficacy of two commercial and indigenous probiotics, Bacillus subtilis and Bacillus licheniformis on growth performance, immuno-physiology and resistance response of juvenile white shrimp (Litopenaeus vannamei). Aquaculture. 2018;496(1):43–9.

35. Blajman JE, Olivero C, Fusari ML, Zimmermann JA, Rossler E, Berisvil AP, et al. Impact of lyophilized Lactobacillus salivarius DSPV 001P administration on growth performance, microbial translocation, and gastrointestinal microbiota of broilers reared under low ambient temperature. Research in veterinary science. 2017;114:388–94.

36. Khan I, Nawaz M, Anjum AA, Ahmad M-u-D, Mehmood A, Rabbani M, et al. Effect of Indigenous Probiotics on Gut Morphology and Intestinal Absorption Capacity in Broiler Chicken Challenged with Salmonella enteritidis. Pakistan Journal of Zoology. 2020;52(5):1825.

37. Shokryazdan P, Faseleh Jahromi M, Liang JB, Ramasamy K, Sieo CC, Ho YW. Effects of a Lactobacillus salivarius mixture on performance, intestinal health and serum lipids of broiler chickens. PLoS one. 2017;12(5):e0175959.

38. Mountzouris K, Tsitrsikos P, Palamidi I, Arvaniti A, Mohnl M, Schatzmayr G, et al. Effects of probiotic inclusion levels in broiler nutrition on growth performance, nutrient digestibility, plasma immunoglobulins, and cecal microflora composition. Poultry science. 2010;89(1):58–67.

39. Revolledo L, Ferreira C, Ferreira A. Prevention of Salmonella Typhimurium colonization and organ invasion by combination treatment in broiler chicks. Poultry science. 2009;88(4):734–43.

40. Bertelloni F, Tosi G, Massi P, Fiorentini L, Parigi M, Cerri D, et al. Some pathogenic characters of paratyphoid Salmonella enterica strains isolated from poultry. Asian Pacific journal of tropical medicine. 2017;10(12):1161–6.

41. Abdel-Hafeez HM, Saleh ES, Tawfeek SS, Youssef IM, Abdel-Daim AS. Effects of probiotic, prebiotic, and synbiotic with and without feed restriction on performance, hematological indices and carcass characteristics of broiler chickens. Asian-Australasian journal of animal sciences. 2017;30(5):672.

42. Awad W, Aschenbach J, Khayal B, Hess C, Hess M. Intestinal epithelial responses to Salmonella enterica serovar Enteritidis: effects on intestinal permeability and ion transport. Poultry science. 2012;91(11):2949–57.

43. Harimurti S, editor Effect of indigenous lactic acid bacteria probiotics on broiler performance. International Seminar on Tropical Animal Production (ISTAP); 2010.

44. Cao G, Zeng X, Chen A, Zhou L, Zhang L, Xiao Y, et al. Effects of a probiotic, Enterococcus faecium, on growth performance, intestinal morphology, immune response, and cecal microflora in broiler chickens challenged with Escherichia coli K88. Poultry science. 2013;92(11):2949–55.

45. Aboshady HM, Stear M, Johansson A, Jonas E, Bambou J-C. Immunoglobulins as biomarkers for gastrointestinal nematodes resistance in small ruminants: A systematic review. Scientific reports. 2020;10(1):1–14.

46. Salim H, Kang H, Akter N, Kim D, Kim J, Kim M, et al. Supplementation of direct-fed microbials as an alternative to antibiotic on growth performance, immune response, cecal microbial population, and ileal morphology of broiler chickens. Poultry science. 2013;92(8):2084–90.

47. Chen C, Li J, Zhang H, Xie Y, Xiong L, Liu H, et al. Effects of a probiotic on the growth performance, intestinal flora, and immune function of chicks infected with Salmonella pullorum. Poultry Science. 2020;99(11):5316–23.

48. Nayebpor M, Farhomand P, Hashemi AJJ. Effects of different levels of direct fed microbial (Primalac) on growth performance and humoral immune response in broiler chickens. Advances in Animal and Veterinary Sciences. 2007;6(11):1308–13.

49. Rahimi M. Effects of probiotic supplementation on performance and humoral immune response of broiler chickens. Book of proceedings, 2nd Mediterranean Summit of WPSA. 2009:67-9.

50. Scharek L, Guth J, Filter M, Schmidt MF. Impact of the probiotic bacteria Enterococcus faecium NCIMB 10415 (SF68) and Bacillus cereus var. toyoi NCIMB 40112 on the development of serum IgG and faecal IgA of sows and their piglets. Archives of animal nutrition. 2007;61(4):223–34.

51. Vilà B, Fontgibell A, Badiola I, Esteve-Garcia E, Jiménez G, Castillo M, et al. Reduction of Salmonella enterica var. Enteritidis colonization and invasion by Bacillus cereus var. toyoi inclusion in poultry feeds. Poultry Science. 2009;88(5):975–9.

52. Zhang D, Li R, Li J. Lactobacillus reuteri ATCC 55730 and L22 display probiotic potential in vitro and protect against Salmonella-induced pullorum disease in a chick model of infection. Research in veterinary science. 2012;93(1):366–73.

53. Luoma A, Markazi A, Shanmugasundaram R, Murugesan G, Mohnl M, Selvaraj R. Effect of synbiotic supplementation on layer production and cecal Salmonella load during a Salmonella challenge. Poultry science. 2017;96(12):4208–16.

54. Levkut M, Revajová V, Lauková A, Ševčíková Z, Spišáková V, Faixová Z, et al. Leukocytic responses and intestinal mucin dynamics of broilers protected with Enterococcus faecium EF55 and challenged with Salmonella Enteritidis. Research in veterinary science. 2012;93(1):195–201.

55. Nakphaichit M, Sobanbua S, Siemuang S, Vongsangnak W, Nakayama J, Nitisinprasert S. Protective effect of Lactobacillus reuteri KUB-AC5 against Salmonella Enteritidis challenge in chickens. Benef Microbes. 2019;10(1):43–54.

56. Starke I, Zentek J, Vahjen W. Effects of the probiotic Enterococcus faecium NCIMB 10415 on selected lactic acid bacteria and enterobacteria in co-culture. Beneficial microbes. 2015;6(3):345–52.

57. Vahjen W, Jadamus A, Simon O. Influence of a probiotic Enterococcus faecium strain on selected bacterial groups in the small intestine of growing turkey poults. Archives of animal nutrition. 2002;56(6):419–29.

58. Yu B, Liu J, Chiou M, Hsu Y, Chiou P. The effects of probiotic Lactobacillus reuteri Pg4 strain on intestinal characteristics and performance in broilers. Asian-australasian journal of animal sciences. 2007;20(8):1243–51.

59. Smialek M, Burchardt S, Koncicki A. The influence of probiotic supplementation in broiler chickens on population and carcass contamination with Campylobacter spp.-Field study. Research in veterinary science. 2018;118:312–6.

60. Stephenson DP, Moore RJ, Allison GE. Lactobacillus strain ecology and persistence within broiler chickens fed different diets: identification of persistent strains. Applied Environmental Microbiology. 2010;76(19):6494–503.

